# Chromosome Evolution of Octoploid Strawberry

**DOI:** 10.1101/861526

**Authors:** Michael A. Hardigan, Mitchell J. Feldmann, Anne Lorant, Randi Famula, Charlotte Acharya, Glenn Cole, Patrick P. Edger, Steven J. Knapp

## Abstract

The allo-octoploid cultivated strawberry (*Fragaria* × *ananassa*) originated through a combination of polyploid and homoploid hybridization, domestication of an interspecific hybrid lineage, and continued admixture of wild species over the last 300 years. While genes appear to flow freely between the octoploid progenitors, the genome structures and diversity of the octoploid species remain poorly understood. The complexity and absence of an octoploid genome frustrated early efforts to study chromosome evolution, resolve subgenomic structure, and develop a single coherent linkage group nomenclature. Here, we show that octoploid *Fragaria* species harbor millions of subgenome-specific DNA variants. Their diversity was sufficient to distinguish duplicated (homoeologous and paralogous) DNA sequences and develop 50K and 850K SNP genotyping arrays populated with co-dominant, disomic SNP markers distributed throughout the octoploid genome. Whole-genome shotgun genotyping of an interspecific segregating population yielded 1.9M genetically mapped subgenome variants in 5,521 haploblocks spanning 3,394 cM in *F. chiloensis* subsp. *lucida*, and 1.6M genetically mapped subgenome variants in 3,179 haploblocks spanning 2,017 cM in *F*. × *ananassa*. These studies provide a dense genomic framework of subgenome-specific DNA markers for seamlessly cross-referencing genetic and physical mapping information, and unifying existing chromosome nomenclatures. Through comparative genetic mapping, we show that the genomes of geographically diverse wild octoploids are effectively diploidized and completely collinear. The preservation of genome structure among allo-octoploid taxa is a critical factor in the unique history of garden strawberry, where unimpeded gene flow supported both its origin and domestication through repeated cycles of interspecific hybridization.

## Introduction

Interspecific homoploid hybridization and polyploidy-inducing hybrid events have been creative forces in plant genome evolution and speciation, acting as catalysts for *de novo* reorganization of chromosome structure (Alix et al., 2017; Jiao et al., 2011; Mandáková et al., 2019; McKain et al., 2016; Soltis et al., 2014b, 2014a, 2016; Vallejo-Marín et al., 2015; Wendel et al., 2016; Yakimowski and Rieseberg, 2014). The cultivated strawberry (*Fragaria* × *ananassa* Duchesne ex Rozier) is unique among domesticated crop species because it arose through both processes. The chromosomes of octoploid garden strawberry (2n = 8x = 56) evolved through a combination of ancient polyploidy, and repeated homoploid hybridization in the last three centuries (Darrow, 1966; Duchesne, 1766). The presence of duplicated (homoeologous) chromosomes in plants frequently leads to meiotic anomalies and associated chromosomal rearrangements, e.g., translocations and inversions, that reduce or eliminate gene flow between the donors and their polyploid offspring (Alix et al., 2017; Latta et al., 2019; Soltis et al., 2014b, 2014a). Similarly, meiotic mispairing in interspecific homoploid hybrids can lead to rearranged offspring chromosomes that differ from the chromosomes of one or both parents, resulting in reproductive barriers and hybrid speciation, as has been widely documented in sunflower (*Helianthus*) and other plants (Abbott et al., 2010; Barb et al., 2014; Burke et al., 2004; Rieseberg, 1997; Yakimowski and Rieseberg, 2014). However, reproductive barriers among octoploid *Fragaria* taxa remain essentially nonexistent, fueling the recurrence of interspecific homoploid hybridization in the origin, domestication, and modern-day breeding of *F.* × *ananassa*.

The modern *F.* × *ananassa* lineage traces its origin to extinct cultivars developed in western Europe in the 1700s. These cultivars were interspecific hybrids of non-sympatric wild octoploids from the New World: *F. chiloensis* subsp. *chiloensis* from South America and *F. virginiana* subsp. *virginiana* from North America (Darrow, 1966). Repeated introgression of alleles from diverse subspecific ecotypes of *F. virginiana* and *F. chiloensis* defined the later generations, coinciding artificial selection of horticulturally important traits among hybrid descendants in Europe and North America. Modern cultivars have emerged from 250 years of global migration and breeding within this admixed population (Darrow, 1966; Hardigan et al., 2018). Because alleles have been introgressed from up to eight octoploid subspecies, the genomes of modern *F*. × *ananassa* individuals are mosaics of their wild ancestors (Hardigan et al., 2018; Liston et al., 2014). Since the discovery of artificial hybrids at the Gardens of Versaille (Duchesne, 1766), natural interspecific hybrids (*F*. × *ananassa* subsp. *cuneifolia*) were discovered in zones of sympatry between *F. chiloensis* subsp. *pacifica* and *F. virginiana* subsp. *platypetala* in western North America (Hancock Jr and Bringhurst, 1979; Luby et al., 1992; Salamone et al., 2013; Staudt, 1999). Neither cultivated *F*. × *ananassa* or wild *F*. × *ananassa* subsp. *cuneifolia* is reproductively isolated from their octoploid progenitors. Thus, genes appear to flow freely between the wild octoploid progenitors, and between the hybrids and their progenitors. While genomic rearrangements have been identified between homoeologous chromosomes, and relative to diploid species (Tennessen et al., 2014; van Dijk et al., 2014), the apparent absence of reproductive isolation implies that homoploid and polyploid hybridization events have not produced significant chromosome rearrangements among octoploid taxa. We hypothesized that the octoploids carry nearly collinear chromosomes tracing to the most recent common ancestor, despite one million years of evolution which produced multiple recognized species and subspecies.

The octoploid strawberry genome has been described as “notoriously complex” and an “extreme example of difficulty” for study (Folta and Davis, 2006; Hirakawa et al., 2014; Hirsch and Buell, 2013; Koskela et al., 2016). While GBS, GWAS, and NGS-reliant applications are relatively straightforward in organisms with well-characterized reference genomes, such approaches were previously difficult or intractable in octoploid strawberry (Bassil et al., 2015; Liston et al., 2014; Tennessen et al., 2014; Vining et al., 2017). Genetic studies in octoploid strawberry previously relied on the genome of woodland strawberry (*F. vesca*), an extant relative to one of four diploid subgenomes contained in *F*. × *ananassa* (Edger et al., 2017; Shulaev et al., 2011). For nearly a decade, the *F. vesca* genome was the only framework available for DNA variant discovery, gene discovery, genetic mapping, and genome-wide association studies in octoploid strawberry (Bassil et al., 2015; Davik et al., 2015; Pincot et al., 2018; Tennessen et al., 2014; Vining et al., 2017). The development of a chromosome-scale reference genome for *F*. × *ananassa* (Edger et al., 2019b) provided the physical framework needed to overcome previous barriers, and explore the organization and evolution of its progenitor genomes.

Here, we report the first study of chromosome evolution and genome structure in octoploid *Fragaria* using an octoploid genome-guided approach to DNA variant discovery and comparative mapping. We demonstrate the ability to differentiate duplicated (homoeologous) octoploid sequences using both NGS and array-based genotyping technologies when applied in conjunction with an octoploid reference genome. In doing so, we overcome a long-standing technical hurdle that has impeded efforts to study strawberry subgenome diversity and chromosome evolution. We estimated strawberry subgenomic diversity by whole-genome shotgun (WGS) sequencing of 93 genealogically and phylogenetically diverse *F.* × *ananassa, F. chiloensis*, and *F. virginiana* individuals. The frequency of unique WGS sequence alignment to the octoploid strawberry genome was characteristic of many diploid plant species (Hamilton and Buell, 2012; Lee and Schatz, 2012; Schatz et al., 2012; Treangen and Salzberg, 2012), and permitted the identification of millions of subgenome-specific DNA variants, effectively distinguishing homologous and homoeologous DNA sequences on every chromosome. Using the genetic diversity of *F*. × *ananassa*, we developed publicly available 50K and 850K SNP arrays populated with subgenome anchored marker probes for octoploid genetic mapping and forward genetic studies. We then performed high-density genetic mapping of five octoploids representing *F.* × *ananassa* and both its progenitor species using a combination of WGS-based and array-based genotyping. Telomere-to-telomere genetic mapping of nearly every chromosome was enabled by the conserved disomic segregation observed in populations derived from wild species and *F.* × *ananassa*, underscoring the effective diploidization and meiotic stability of octoploid *Fragaria*. Comparative mapping of *F.* × *ananassa* and multiple subspecies of *F. chiloensis* and *F. virginiana* revealed the genome structures of the cultivated reference genotype (Camarosa) and its progenitor species were nearly identical.

The collinear and diploidized genomes of *F.* × *ananassa* and its progenitors support octoploid *Fragaria* as an evolutionary clade which achieved a relatively high degree of genome stability prior to the speciation and sub-speciation of *F. chiloensis* and *F. virginiana*. Strawberry’s interspecific origin followed by successive hybridization throughout domestication is an unusual improvement pathway typically associated with perennial tree fruits, and frequently contributes to reproductive incompatibility or sterility in wide species hybrids (Hughes et al., 2007; Ladizinsky, 1985; Meyer and Purugganan, 2013; Miller and Gross, 2011). The preservation of genome structure among diverse octoploid *Fragaria* species and subspecies was likely essential to the unique history of the *F.* × *ananassa* lineage, which has undergone repeated cycles of homoploid hybridization without the formation of reproductive barriers or loss of fertility.

## Results & Discussion

### Subgenomic Diversity of Octoploid Fragaria

We performed the first deep exploration of the homoeologous sequence diversity of octoploid *Fragaria* using the Camarosa v1.0 octoploid reference genome (Edger et al., 2019b) and a diversity panel of 93 strawberry individuals, including 47 *F.* × *ananassa* samples, 24 *F. chiloensis* samples, and 22 *F. virginiana* samples (Table S1). By incorporating subgenome specificity at the assembly level, previous barriers to copy-specific sequence alignment caused by strawberry’s octoploid ancestral homology posed a less significant obstacle than repetitive DNA elements for diploid genomes such as maize (Hamilton and Buell, 2012; Lee and Schatz, 2012; Schatz et al., 2012; Treangen and Salzberg, 2012). The fraction of uniquely aligning (MapQ > 0) PE150 sequences averaged 83.2%, and 90.9% of Camarosa PE250 sequences aligned uniquely (Figure S1), allowing comprehensive coverage and analysis of subgenomic diversity. Using genotype calling software FreeBayes and a series of hard-filters targeting unique sequence alignments, we identified 41.8M subgenomic SNP and indel mutations in *F.* × *ananassa* and its wild progenitors.

*F.* × *ananassa* has been described as “genetically narrow” due to the small number of founders in the pedigrees of modern cultivars (Dale and Sjulin, 1990; Sjulin and Dale, 1987; Stegmeir et al., 2010). Despite a small effective population size, our analyses show that massive genetic diversity has been preserved in *F.* × *ananassa*, with negligible difference between wild species and domesticated germplasm. The subgenome nucleotide diversity (π) of *F.* × *ananassa* (π = 5.857 x 10-3) was equivalent to wild progenitors *F. chiloensis* (π = 5.854 x 10-3) and *F. virginiana* (π = 5.954 x 10-3), and comparable to the sequence diversity of *Zea mays* landraces (π = 4.9 x 10-3) and wild *Zea mays spp. parviglumis* progenitors (π = 5.9 x 10-3) (Hufford et al., 2012). Correlations of *F.* × *ananassa, F. chiloensis*, and F. *virginiana* diversity across the 28 octoploid chromosomes ranged from 0.93-0.97, showing the magnitude and distribution of genomic diversity are broadly conserved among octoploid taxa. This suggested that *F.* × *ananassa* was not strongly bottlenecked by domestication, or that its domestication bottleneck was mitigated by continued introgression of allelic diversity from wild subspecies (Darrow, 1966). We found that variance in the distribution of octoploid nucleotide diversity was influenced more significantly by subgenome ancestry than domestication and breeding (Table S2). The diploid *F. vesca* subgenome, dominant with respect to gene abundance and expression (Edger et al., 2019b), contained the least diverse homoeolog of every ancestral chromosome, while subgenomes derived from the ancestors of the extant Asian species (*F. iinumae* and *F. nipponica*) contained greater genetic diversity (Table S2). These differences show that subgenome dominance supports distinct levels of purifying selection and genetic diversity on different chromosomes of octoploid strawberry.

Because *F.* × *ananassa* was domesticated as an interspecific hybrid, individual performance is assumed to benefit from allelic diversity between *F. chiloensis* and *F. virginiana*. To support comparisons of strawberry heterozygosity with previously studied polyploid species, we estimated individual heterozygosity based on the genomic frequency of heterozygous nucleotides (nts) (potato metric) (Hardigan et al., 2017), and the frequency of heterozygosity at GBS-derived polymorphic sites (blueberry and cotton metric) (de Bem Oliveira et al., 2019; Page et al., 2013). Strawberry genomic heterozygosity ranged from 0.02-0.80% and averaged 0.46% genomic nts. This translated to an average of 11.1% heterozygosity at polymorphic marker sites. The most heterozygous octoploid genomes, White Carolina (1700s – 0.80% nts), PI551736 (Peruvian landrace – 0.72% nts), Jucunda (mid-1800s – 0.70% nts), and Ettersberg 121 (early 1900s – 0.66% nts), were heirloom *F.* × *ananassa* types with more recent *F. virginiana* x *F. chiloensis* parentage. The average subgenomic heterozygosity of octoploid strawberry (0.46% nts) was below diploid potato (1.05% nts) and tetraploid potato (2.73% nts) (Hardigan et al., 2017). The average heterozygosity of octoploid strawberry at GBS-derived polymorphic sites was below the allo-tetraploid cotton A-genome (13% marker sites), above the cotton D-genome (<1% marker sites) (Page et al., 2013), and below autotetraploid blueberry (32.4% marker sites) (de Bem Oliveira et al., 2019). Due to the presence of four ancestral homoeologs, traditional models of “fixed heterozygosity” applied to allopolyploid species (Comai, 2005; Obbard et al., 2006) assume an octoploid functional heterozygosity four-fold greater than subgenomic estimates. Under this model, recent *F. virginiana* x *F. chiloensis* hybrids such as White Carolina and Jucunda could be regarded as similarly heterozygous to autopolyploid species such as potato. However, assembly of the allo-octoploid strawberry genome uncovered rampant gene silencing, gene loss, and rearrangements relative to diploid ancestors (Edger et al., 2019b), eroding the conservation of ancestral allele function. The frequency of unique sequence alignment (Figure S1) and unbroken distribution of subgenomic variant detection (Figure S2) in our analysis underscore the extensive divergence of the four subgenomes. Thus, traditional polyploid allele dosage models assuming genome-wide fixed heterozygosity may be of limited usefulness for strawberry.

### Recombination Breakpoint Mapping of Octoploid Strawberry

We used WGS sequence analysis and recombination breakpoint mapping of an octoploid strawberry population to explore the breadth of disomic variation as an indicator of bivalent pairing during meiosis. Several cytogenetic and DNA marker studies have proposed the occurrence of polysomy in strawberry (Fedorova, 1946; Lerceteau-Köhler et al., 2003; Senanayake and Bringhurst, 1967), while others suggest that octoploids are mainly disomic (Arulsekar and Bringhurst, 1981; Bringhurst, 1990; Byrne and Jelenkovic, 1976; Kunihisa et al., 2005). We performed low-coverage (4-8x) sequencing and subgenomic variant calling in a population (n = 189) derived from a cross between the *F.* × *ananassa* reference genotype Camarosa and the beach strawberry (*F. chiloensis* subsp. *lucida*) ecotype Del Norte. These parents were selected to provide a dense comparison of profiles of mappable diploid DNA variation in a natural octoploid and an artificial hybrid (reference genotype). Variant calling against the Camarosa v1.0 genome identified 3.7M subgenomic SNPs and indels inherited from 1.6M Camarosa heterozygous sites (AB × AA), 1.9M Del Norte heterozygous sites (AA × AB), and 0.2M co-heterozygous sites (AB × AB). We used the high-density variant data to perform haplotype mapping based on recombination breakpoint prediction, and evaluated segregation ratios of parental alleles across the 28 octoploid chromosomes.

We bypassed the computational demand of analyzing pairwise linkage across millions of DNA variants with missing data and genotyping errors by implementing the haplotype calling approach proposed by Huang et al. (2009) and Marand et al. (2017). Our approach performed a sliding-window analysis to predict crossover events, then estimated the consensus of co-segregating DNA variation between recombination breakpoints to reconstruct the representative genotypes of each haploblock, which were mapped as unique genetic markers. Using this approach, we mapped 1.9M Del Norte variants in 5,521 haploblocks spanning 3,393.86 cM, and 1.6M Camarosa variants in 3,179 haploblocks spanning 2,016.95 cM (Dataset S1). The paternal beach strawberry (Del Norte) map produced telomere-to-telomere coverage of the 28 octoploid chromosomes (Figure 1), providing the most comprehensive genetic map of an octoploid *Fragaria* genome to-date. The complete mapping of the extant homoeologs for all seven ancestral *Fragaria* chromosomes in Del Norte, and analysis of chromosome-wide segregation distortion (Figure 2), showed that disomic recombination is ubiquitous in the genome of *F. chiloensis*. By contrast, less than 50% of the Camarosa genome could be mapped on chromosomes 1-1, 1-2, 2-4, 3-3, 5-2, 6-2, 6-3, 6-4, and 7-3 (Figure S3). We analyzed Camarosa heterozygosity and segregation distortion to determine whether the inability to map large segments of the genome was the result of polysomic recombination in *F.* × *ananassa*. This uncovered a near total loss of polymorphism in the unmapped regions of Camarosa (Figure 2), showing that incomplete mapping of *F.* × *ananassa* resulted from depletion of heterozygosity in the hybrid genome, not polysomy. Sargent et al. (2012) previously reported extensive regions of homozygosity that affected mapping of *F.* × *ananassa*. Artificial selection pressure in commercially bred hybrids almost certainly accounts for the lower subgenomic heterozygosity of Camarosa relative to Del Norte, which does not support a critical role for genome-wide interspecific heterozygosity in driving cultivar performance.

**Figure 1.**
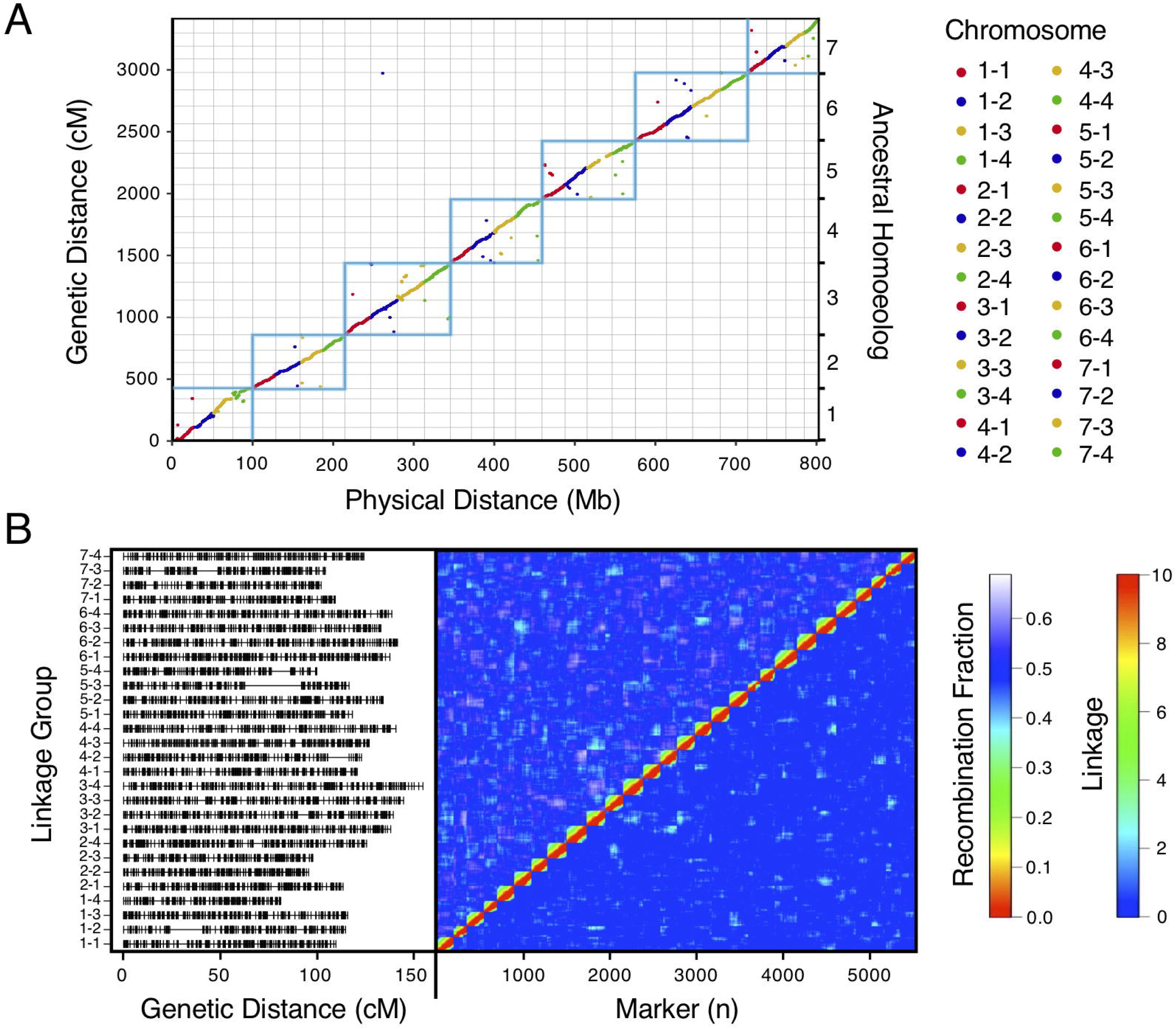
High-density haplotype map of a California beach strawberry (*F. chiloensis* subsp. *lucida*) genome. (A) Del Norte genetic map distances plotted against the Camarosa v1.0 physical genome. Box outlines indicate groups of ancestral chromosome homoeologs. (B) Del Norte linkage groups plotted with corresponding chromosome heatmap of pairwise recombination fractions (upper diagonal) and pairwise linkage (lower diagonal).

**Figure 2.**
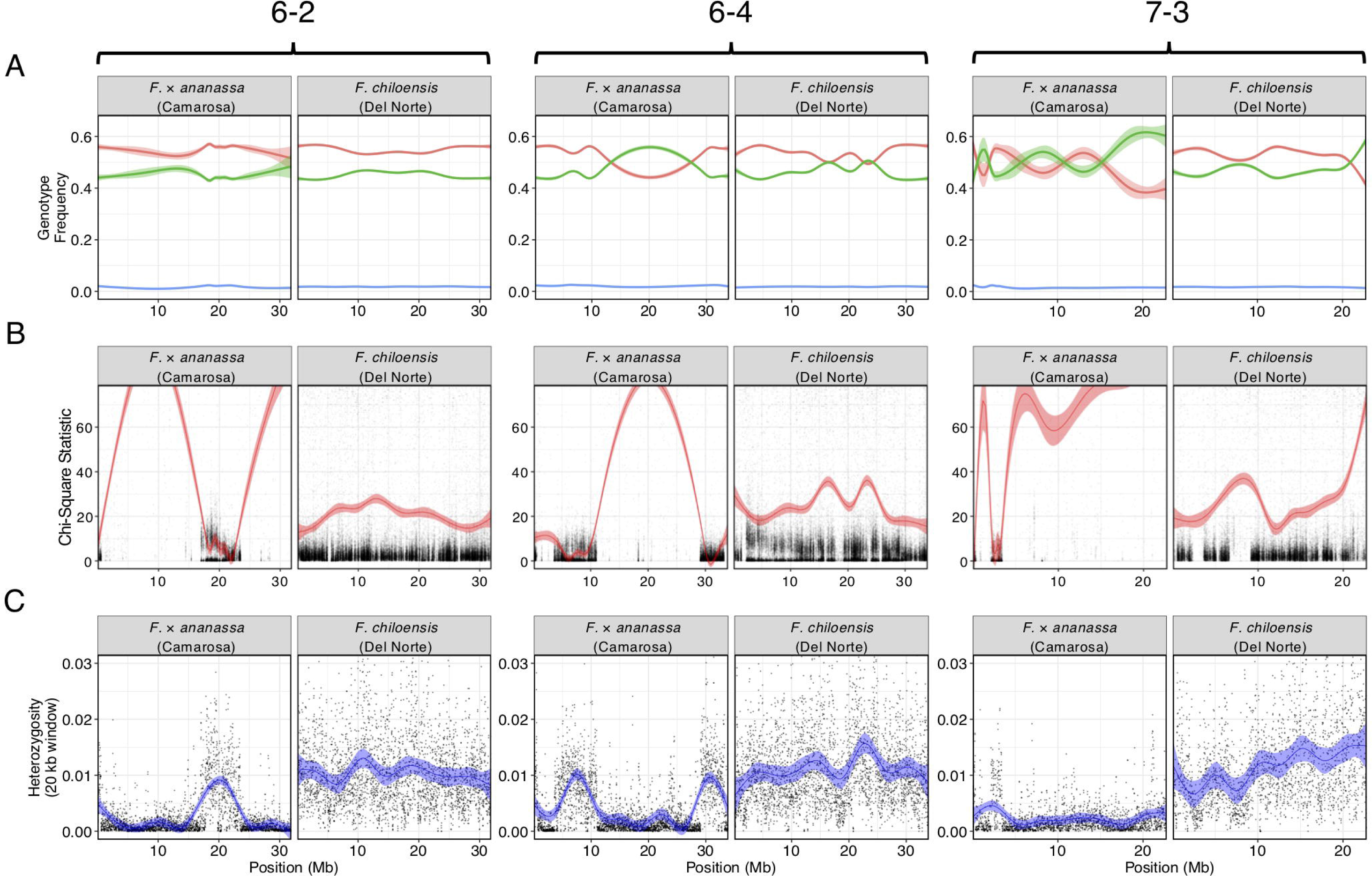
Comparison of genomic heterozygosity and diploid segregation distortion in *F.* × *ananassa* (Camarosa) and *F. chiloensis* (Del Norte) on three chromosomes (6-2, 6-4, 7-3). (A) Frequency of AA (green), AB (red), and off-target (BB; blue) genotypes for polymorphic markers in the mapping population. (B) Chi-square statistic estimating segregation distortion of polymorphic markers in the mapping population. (C) Heterozygous nucleotide frequency of parent genotypes in 20 kb physical windows.

### 850K Octoploid Screening Array

We designed Affymetrix SNP genotyping arrays populated with subgenome-specific marker probes to enable genetic mapping, genome-wide association studies (GWAS), and genomic prediction in octoploid strawberry. DNA variants were selected for array design from the subgenomic diversity identified in the WGS panel (Figure 3). From the 90M total unfiltered variant sites, we extracted 45M unfiltered variants that segregated in *F.* × *ananassa*. To identify candidate DNA variants for marker design, we selected only biallelic SNPs above a low-diversity threshold (π ≥ 0.05), excluded rare alleles (MAF ≥ 0.05), required a VCF quality score > 20, and excluded sites with > 15% missing data in the diversity panel. These filters yielded 8M subgenomic SNPs segregating within the *F.* × *ananassa* subset of the diversity panel. We obtained 71-nt marker probes by extracting 35-nt sequences flanking each SNP site from the Camarosa v1.0 genome assembly. Marker probes for the 8M high-confidence SNP sites were then filtered to remove candidates that were problematic for array tiling. These included duplicate or near-duplicate probe sequences, probes that inherited ambiguous reference sequences (Ns), probes requiring double-tiling (A/T or C/G alleles), and probes that Affymetrix scored as having low buildability. We retained 6.6M probes that targeted high-confidence *F.* × *ananassa* variants and were acceptable for array tiling.

**Figure 3.**
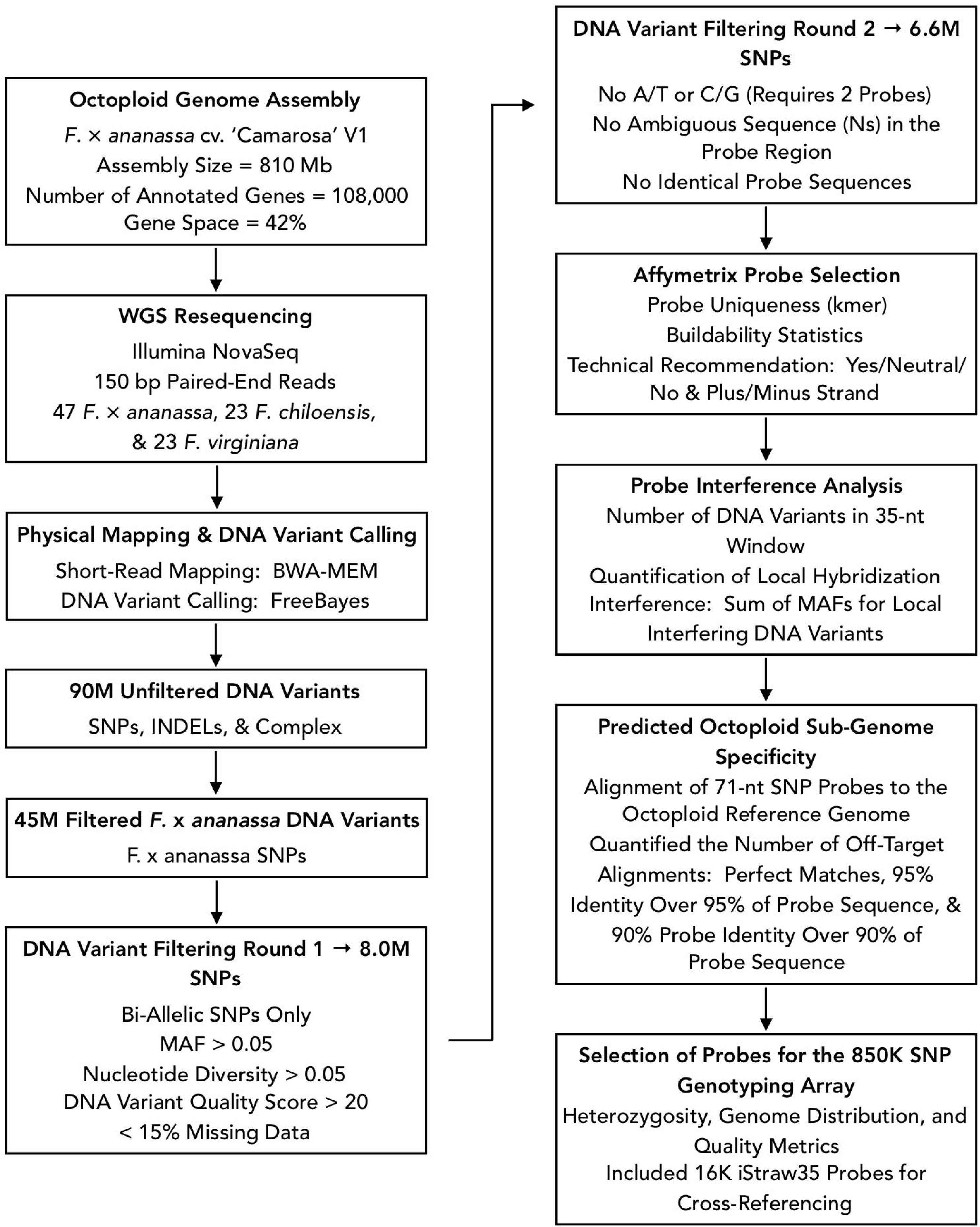
Flowchart of bioinformatic protocols used to select genome-wide sequence variants for design of the 850K SNP screening array.

We applied three selection criteria for determining a subset of 850K marker probes for tiling a screening array: likelihood of probe binding interference by off-target variants, likelihood of off-target (non-single copy) probe binding, and physical genome distribution. The likelihood of probe binding interference was scored as the sum of non-reference allele frequencies for off-target variants in the 35-nt binding region adjacent to the target SNP. The likelihood of off-target probe binding was scored by performing BLAST alignment of the 71-nt probe sequences to the Camarosa v1.0 genome and quantifying the number of off-target alignments with query coverage above 90% and sequence identity above 90%. We then iteratively parsed the Camarosa v1.0 genome using 10 kilobase (kb) non-overlapping physical windows, extracting the best available marker from each window based on probe binding interference and off-target binding likelihoods, until reaching an 850K probe threshold. We reserved 16K positions for legacy markers from the iStraw SNP array (Bassil et al., 2015; Verma et al., 2016) that were polymorphic in a previous strawberry diversity study (Hardigan et al., 2018). The set of 850K probe sequences was submitted to Affymetrix for constructing a screening array.

We genotyped a genetically diverse sample of 384 octoploid strawberry accessions to validate subgenome-specific marker performance on the 850K screening array (Dataset S2). The sample fluorescence files were analyzed with the Axiom Suite in polyploid mode to generate marker clusters. Collectively, 446,644 of 850,000 marker probes produced QC-passing polymorphic genotype clusters showing disomic (allopolyploid) segregation. Among these, 78.3% were classified as “PolyHighResolution” in the Axiom terminology, producing diploid co-dominant genotype clusters (AA, AB, and BB) without detecting off-target allelic variation. Similarly, 18.8% of markers classified as “NoMinorHomozygote” in the Axiom terminology produced dominant genotype clusters in which heterozygotes (AB) clustered with one of the homozygous genotype classes (AA or BB). The remaining 2.9% of the markers showed detection of non-target alleles, and were classified as “OffTargetVariant” markers in the Axiom terminology. The HomRO statistic generated by the Axiom Suite estimates genotype cluster separation, and has been used as a metric to infer octoploid single-copy (i.e. subgenome or paralog specific) probe binding when values exceed 0.3 (Bassil et al., 2015). Based on this threshold, 74% of the QC-passing marker probes on the 850K screening array exhibit single-copy binding, in addition to measuring subgenome-specific DNA variation (Figure S4). The complete set of 446,644 validated probes is made available for public use (Dataset S3).

### 50K Octoploid Production Array

We selected 49,483 polymorphic marker probes from the 850K validated probe set to build a 50K production array (Dataset S4). 5,809 LD-pruned (r^2^ < 0.50) marker probes were retained from the iStraw panel to support cross-referencing of octoploid QTL studies and linkage group nomenclatures. We targeted 2,878 genes based on Camarosa v1.0 functional annotations that indicated R-gene affiliated protein domains (Edger et al., 2019b) or homology to *Fragaria vesca* genes involved in flowering and fruit development expression networks (Hollender et al., 2014; Kang et al., 2013). Candidate genes were pre-allocated up to two markers (within 1 kb) from the screening panel. We next selected a set of the most commonly segregating markers to support genetic mapping. We identified this set by selecting the marker with the highest pairwise diversity (π) in *F.* × *ananassa* across non-overlapping 50 kb physical genome windows. The remainder of the 50K array was populated by iteratively parsing the genome with 50 kb physical windows and selecting random QC-passing markers to provide an unbiased genome distribution. Both the 850K and 50K probe sets provide unbroken, telomere-to-telomere physical coverage of the 28 octoploid strawberry chromosomes (Figure S5). Within the 50K probe set, 53% of the probes were located within genes, and 79% were located within 1 kb of a gene. The 50K probe set was provided to Affymetrix for building the production array.

We screened 1,421 octoploid samples from multiple bi-parental populations and a large diversity panel on the 50K production array. Collectively, 42,081 markers (85%) successfully replicated QC-passing polymorphic genotype clusters when screened in the larger sample group. Of the 7,402 non-replicated markers, only 1% were excluded due to becoming monomorphic (“MonoHighResolution” Axiom class) or from increased missing data (“CallRateBelowThreshold” Axiom class). Sub-clustering and increased dispersion within the AA or BB genotype clusters (“AAvarianceX”, “AAvarianceY”, “BBvarianceX”, “BBvarianceY” Axiom classes) accounted for 20% on non-replicated markers. The Axiom software provided no specific cause for failure for the remaining 79% of non-replicated markers. These results suggest that increasing the size and diversity of a genotyping population may affect the reproducibility of a fraction (15%) of markers on the 50K array, while a majority (85%) are highly reproducible. The fraction of polymorphic co-dominant (“PolyHighResolution”) markers increased from 78% on the 850K screening array to 86% among reproducible markers 50K production array.

### Genetic Mapping of Wild Octoploid Ecotypes

We demonstrated that the 50K SNP array allows dense genetic mapping of heterozygous regions on all 28 chromosomes of *F.* × *ananassa, F. chiloensis*, and *F. virginiana*. Genetic mapping of *F. chiloensis* and *F. virginiana* provided telomere-to-telomere physical representation of the 28 octoploid chromosomes, and near-complete representation within the individual wild maps (Dataset S1). We selected four octoploid parents from two outcrossing bi-parental populations genotyped on the 50K array for mapping. The first population was the Camarosa x Del Norte (*F. chiloensis* subsp. *lucida*) population used for WGS recombination breakpoint mapping (n = 182). The second population was derived from a cross between *F. virginiana* subsp. *virginiana* accession PI552277 (female parent) and *F. virginiana* subsp. *virginiana* accession PI612493 (male parent) (n = 96). The number of SNPs segregating in the octoploid parent genotypes varied considerably (Table S3). Camarosa contained the most segregating markers (9,062), followed by PI552277 (5,575), PI612493 (5,464), and Del Norte (2,368). The larger number of markers segregating in Camarosa relative to the wild parent genotypes reflected the array design strategy, which targeted *F.* × *ananassa* diversity. The unbalanced representation of *F. virginiana* and *F. chiloensis* diversity was not expected, because genome-wide variant calls showed similar profiles of heterozygosity and nucleotide diversity in the wild progenitors, and showed that Del Norte was more heterozygous than Camarosa. The higher level of ascertainment bias against *F. chiloensis* diversity that resulted from probing *F.* × *ananassa* alleles supports previous findings that *F. virginiana* diversity is more prevalent in cultivated hybrids (Hardigan et al., 2018). We mapped heterozygous variant sites of the four parent octoploids using software ONEMAP (Margarido et al., 2007) to generate initial linkage groups and markers orders, and BatchMap (Schiffthaler et al., 2017) for marker re-ordering and genetic distance estimation. Despite the ascertainment bias for domesticated allelic diversity on the 50K array, the relatively unbiased distribution of genomic heterozygosity in wild genotypes (Table S3) provided a more complete representation of the wild octoploid genomes than *F.* × *ananassa* (Camarosa) (Figure 4). Large homozygous regions that produced breaks in the Camarosa WGS haplotype map and 50K array map (chromosomes 1-1, 1-2, 2-4, 3-3, 5-2, 6-2, 6-3, 6-4, and 7-3) were clearly featured in the wild genetic maps (Figure 4). Camarosa contained an average of 11.7 (±6.8) SNPs/megabase (Mb) across 28 chromosomes, with as many as 25.2 SNPs/Mb (1-4) and as few as 0.8 SNPs/Mb (1-3), underscoring the scattered distribution of mappable subgenomic diversity in the commercial hybrid.

**Figure 4.**
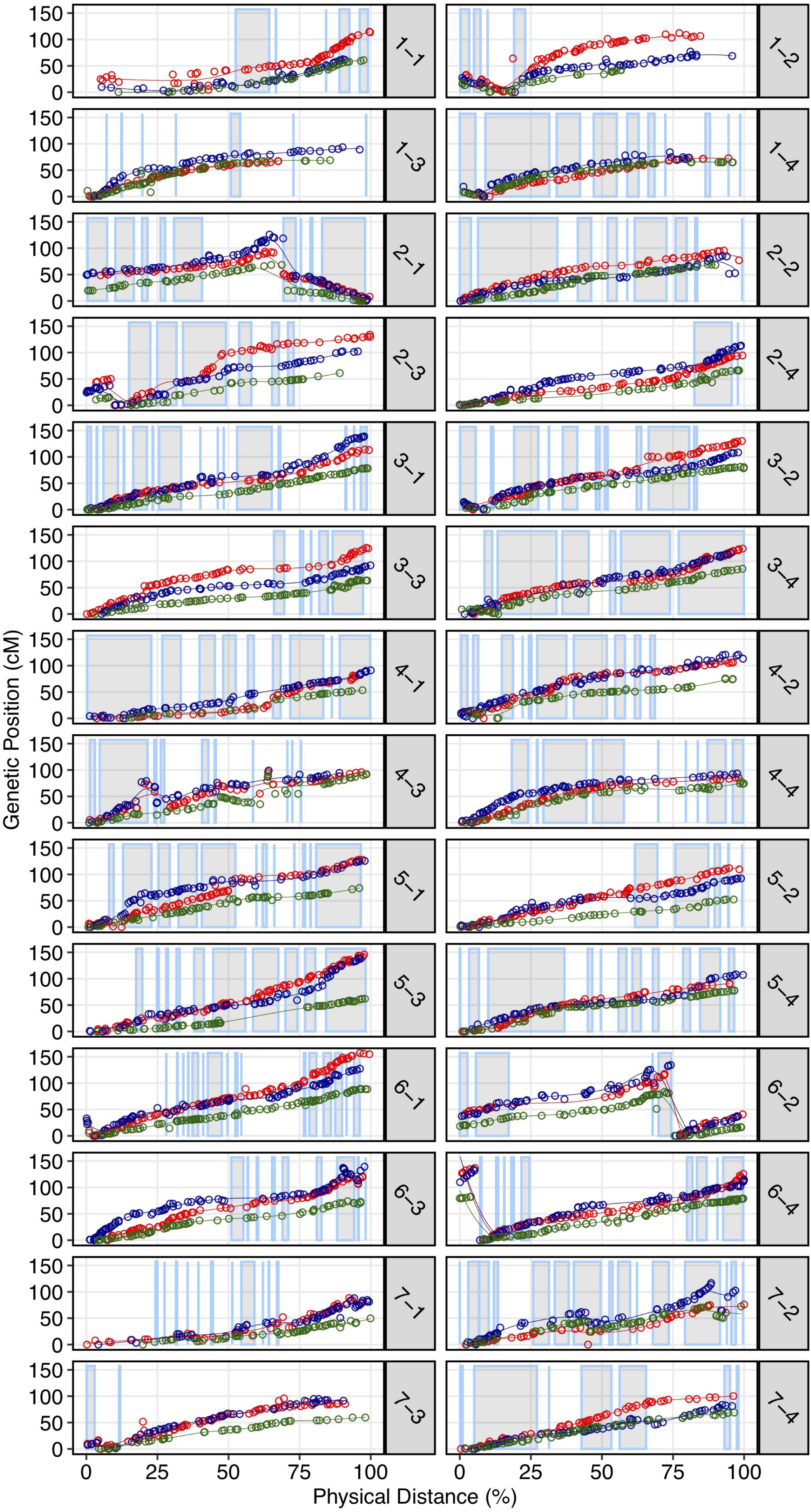
Genetic maps of three wild octoploid strawberry genotypes (PI552277–red; PI612493–blue; Del Norte–green) based on 50K SNP array genotypes plotted against the Camarosa v1.0 physical genome. Grey highlighted chromosome segments indicate contiguous (up to 500 kb) regions of the physical genome represented by the Camarosa 50K SNP array map.

The wild octoploid maps revealed large (Mb+) chromosomal rearrangements relative to the Camarosa v1.0 physical genome on chromosomes 1-2, 1-4, 2-1, 2-3, 6-2, and 6-4. These rearrangements were conserved across the wild species genomes, and supported by corresponding regions represented in the Camarosa genetic map (1-2, 1-4, 2-1) (Figure S3), indicating intra-chromosomal scaffolding errors in the physical reference genome. The fraction of SNPs genetically mapping to non-reference chromosomes ranged from 1.5-1.9% in the four parents, with the highest fraction observed in Camarosa. This indicated minimal inter-chromosomal errors in the physical genome, and minimal inter-chromosomal marker discordance between octoploid progenitor species. Thus, the octoploid genetic maps provided no evidence of chromosome rearrangement among the octoploid species, or relative to *F.* × *ananassa*.

### Genome Structure of Ancestral Octoploids

Previous studies have reported octoploid chromosome rearrangements relative to diploid *Fragaria*, potentially contributing to sex determination (Govindarajulu et al., 2015; Spigler et al., 2008, 2010; Tennessen et al., 2014), and there is phylogenetic evidence of chromosome exchanges among the four ancestral subgenomes (Edger et al., 2019b; Liston et al., 2014). However, there is no evidence for chromosome-scale structural variation between octoploid taxa. It remains unclear to what extent the structural variation of octoploid *Fragaria* reflects initial polyploid ‘genome shock’ associated with a common ancestor, or rather, ongoing mutations contributing to octoploid species diversification. Through comparative mapping, we show that the genomes of diverse octoploid ecotypes contributing to the homoploid hybrid lineage of *F.* × *ananassa* are nearly completely collinear. We constructed genetic maps for two additional wild genotypes, *F. chiloensis* subsp. *pacifica* (SAL3) and *F. virginiana* subsp. *platypetala* (KB3) to capture a more diverse set of octoploid subspecific taxa. We aligned publicly available DNA capture sequences from an *F. chiloensis* subsp. *chiloensis* × *F. chiloensis* subsp. *pacifica* population (GP33 × SAL3, n = 46; Tennessen et al., 2014), and an *F. virginiana* subsp. *platypetala* × *F. virginiana* subsp. *platypetala* population (KB3 × KB11, n = 46; Tennessen et al., 2018) to the Camarosa v1.0 genome assembly, and predicted subgenome variant genotypes using FreeBayes. Genetic mapping of the DNA capture markers followed the protocol used for the 50K SNP datasets. Using the 50K array linkage groups and DNA capture linkage groups (Dataset S1), we performed comparative mapping of four octoploid subspecies – *F. chiloensis* subsp. *lucida* (Del Norte), *F. chiloensis* subsp. *pacifica* (SAL3), *F. virginiana* subsp. *platypetala* (KB3), and *F. virginiana* subsp. *virginiana* (PI552277) – in ALLMAPS based on genetically mapped variant sites anchored to 50-kb physical genome windows in Camarosa v1.0. The chromosomes of the octoploid progenitor subspecies were completely syntenic (Figure 5, Figure S6). Based on these results, large-scale chromosome rearrangements in octoploid *Fragaria* relative to the diploid ancestral genomes would have occurred early on, before the speciation of *F. chiloensis* and *F. virginiana*.

**Figure 5.**
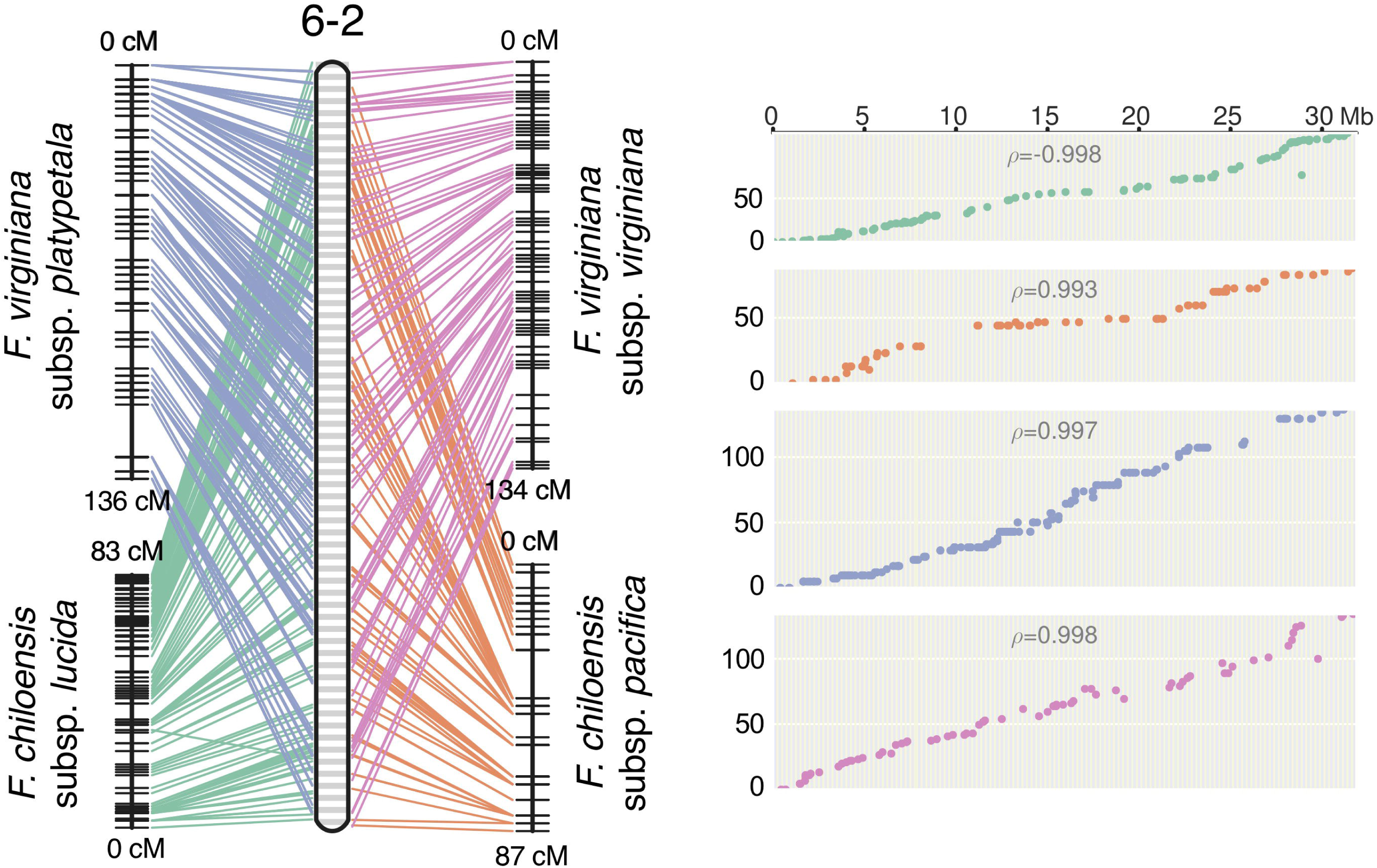
Comparative mapping of four wild octoploid *Fragaria* subspecies (*F. chiloensis* subsp. *lucida, F. chiloensis* subsp. *pacifica, F. virginiana* subsp. *platypetala, F. virginiana* subsp. *virginiana*) on chromosome 6-2.

### Unification of Octoploid Chromosome Nomenclatures

Previous octoploid genetic mapping studies relied on a variety of DNA marker technologies including early PCR-based assays (microsatellites, AFLPs), and later, technical ploidy reduction of sequence variants called against the diploid *F. vesca* genome based on DNA capture, GBS, or WGS-derived array probes (Bassil et al., 2015; Rousseau-Gueutin et al., 2008; Spigler et al., 2008, 2010; Tennessen et al., 2014, 2018; Vining et al., 2017). This diversity of marker genotyping strategies without information linking to the *F.* × *ananassa* physical genome has caused a proliferation of disconnected strawberry chromosome nomenclatures that may not accurately reflect the phylogenetic origins of its respective subgenomes. The Camarosa v1.0 reference genome provides an anchoring point for unifying the existing octoploid nomenclatures. We aligned all historic *Fragaria* microsatellite markers in the Rosacea Genomic Database (GDR) to the Camarosa v1.0 genome and anchored the Spigler et al. (2010) nomenclature to the Camarosa physical genome, which provided the corresponding linkage groups for anchoring the Tennessen et al. (2014) nomenclature to Camarosa v1.0. We then utilized legacy iStraw probes retained on the 50K array to link the Sargent and van Dijk chromosome nomenclatures (Sargent et al., 2016; van Dijk et al., 2014) to the Camarosa v1.0 genome, which was scaffolded using the map published by Davik et al. (2015). In total, five of the most widely cited octoploid strawberry chromosome nomenclatures were unified in relation to the physical genome (Table 1). The existing octoploid nomenclatures each contained subgenome assignments that were incongruent with ancestral chromosomal origins determined by phylogenetic analysis of the physical genome (Edger et al., 2019b, 2019a), particularly with respect to the non-*vesca* and non-*iinumae* subgenomes. The unmasking of octoploid homoeologous chromosome lineages (Edger et al., 2019b, 2019a), and construction of genetic maps showing complete collinearity among ancestral species, provide the foundation for a unified octoploid nomenclature reflecting the phylogenetic origins of its subgenomes.

**Table 1.**
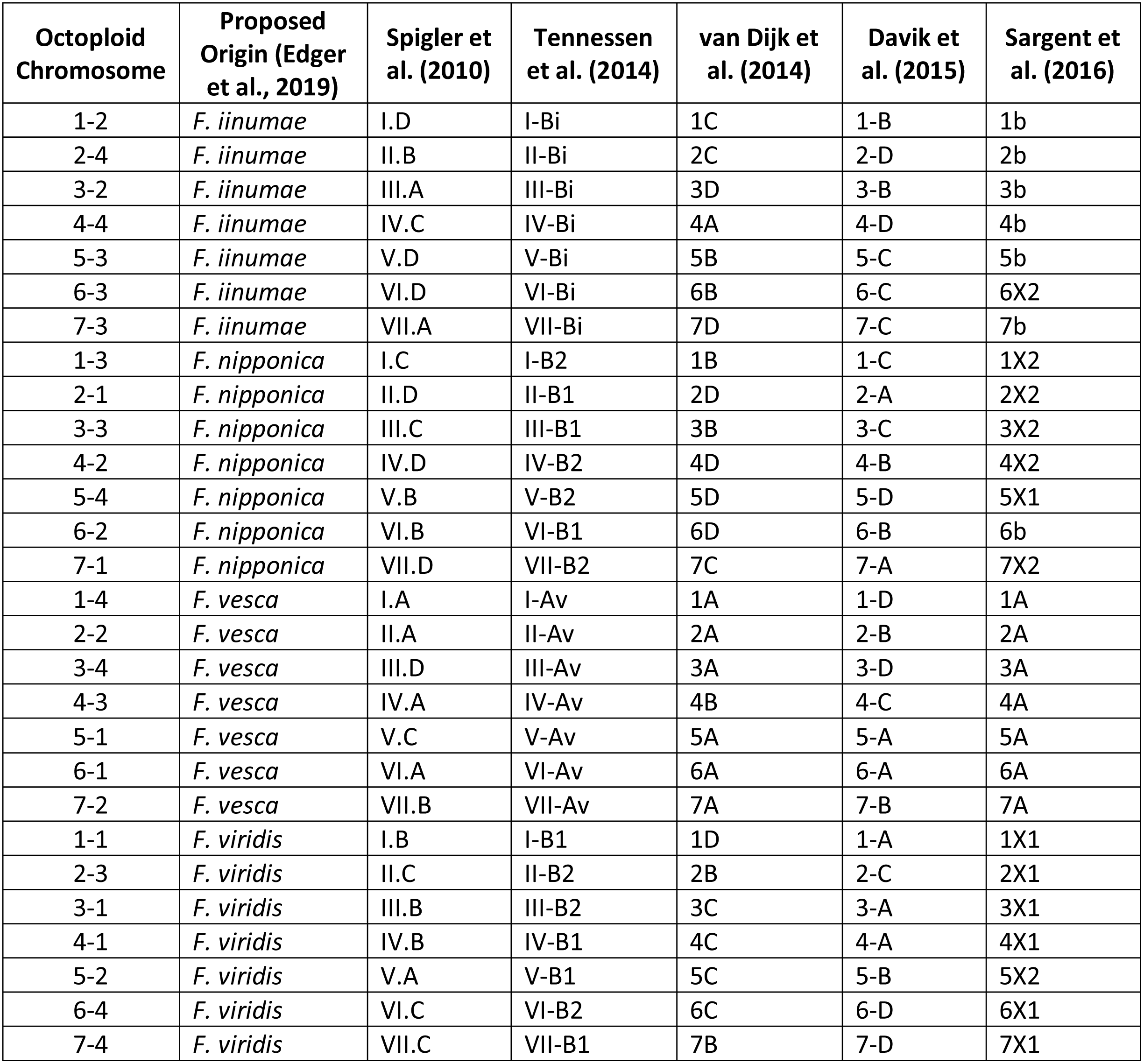
Published octoploid strawberry linkage group nomenclatures anchored to corresponding physical chromosomes in the Camarosa v1.0 reference genome.

## Conclusion

Using the first octoploid genome-guided approach to subgenomic (diploid) variant discovery, we have demonstrated that the genomes of the octoploid progenitors of *F*. × *ananassa* are highly collinear and diploidized (Figure 5, Figure S6). Octoploid *Fragaria* taxa do not follow the common polyploid rule book for chromosome rearrangement (Alix et al., 2017; Chester et al., 2012; Cifuentes et al., 2010; Gaeta et al., 2007; Gaeta and Pires, 2010; Leitch and Leitch, 2008; Ramsey and Schemske, 2002; Renny-Byfield and Wendel, 2014; Wendel et al., 2016), but instead exhibit incredible karyotypic stability across biogeographically diverse subspecies. Strikingly, we did not observe any large-scale (Mb+) structural rearrangements (e.g., translocations or inversions) in the genomes of *F. chiloensis, F. virginiana*, or *F. × ananassa* (Figure 4). The broad conservation of chromosome structure across diverse progenitor taxa partly explains the absence of reproductive barriers and ease of gene flow among currently defined species and the domesticated hybrid lineage. In this regard, octoploid *Fragaria* species appear to be part of a minority among polyploid plants; however, similar examples of karyotypic stability have been described in monocots and dicots (Sun et al., 2017; VanBuren et al., 2019). Because of the ubiquity of polyploidy in angiosperms and the diversity of chromosome-restructuring outcomes along the pathway to diploidization, universal rules do not necessarily apply (Cifuentes et al., 2010; Le Comber et al., 2010; Renny-Byfield and Wendel, 2014; Wendel et al., 2016). The remarkable karyotypic stability and absence of chromosome rearrangements among octoploid *Fragaria* taxa are indicative of regular diploid meiotic behavior and suggest that homoeologous recombination has failed to disrupt the ancestral octoploid karyotype, which has been preserved over 0.4-2.1 M years of chromosome evolution and taxonomic diversification (Tennessen et al., 2014). The unique history of strawberry as a crop lineage, including its origin as an interspecific hybrid and frequent use of interspecific hybridization throughout domestication, was almost certainly supported by an uncommon stability of ancestral genome structure, and resulting ease of gene flow across octoploid genetic backgrounds.

The global importance and rapid commercial success of *F*. × *ananassa* over the last 250 years has been attributed to the interspecific homoploid hybrid component of heterosis (Rho et al., 2012; Shaw, 1995; Spangelo et al., 1971; Stegmeir et al., 2010). While some anecdotal evidence for heterosis exists in *F*. × *ananassa*, quantitative evidence is limited (Rho et al., 2012; Shaw, 1995; Spangelo et al., 1971; Stegmeir et al., 2010). Heterosis is an oft-cited advantage of polyploidy, where genomic heterozygosity is preserved by strict subgenomic recombination, the so-called “fixed heterosis” component of heterosis in allopolyploids (Comai, 2005). The *F*. × *ananassa* genome is riddled with ancient homoeologous exchanges (Edger et al., 2019b), a hallmark of inter-subgenomic recombination in early generations. Neverthless, the genomes of present-day octoploid taxa appear to be highly diploidized. We observed disomic inheritance of DNA variants across the genomes of the octoploids in the present study, and similar ranges of subgenomic heterozygosity for wild individuals and commercial hybrids. The success of *F*. × *ananassa* should not be solely attributed to “fixed heterosis” because neither octoploid progenitor species, which share the effects of fixed heterosis and show similar subgenomic heterozygosity, was commercially successful before the hybrid (Darrow, 1966; Finn et al., 2013). We hypothesize that interspecific complementation, a broader pool of potentially adaptive alleles, and masking of deleterious mutations could be more important than fixed heterosis in *F*. × *ananassa* (Alix et al., 2017; Comai, 2005).

We have shown that the purported complexity and previous intractability of octoploid strawberry genomics were largely associated with the technical challenge of distinguishing subgenome level variation from the broader pool of ancestral sequence homology. The use of an allo-octoploid reference genome addressed this problem by allowing variant calling based on unique sequence alignments to the respective subgenomes. While local subgenome homology could remain an issue, we identified a nearly continuous distribution of subgenome-specific variation spanning the octoploid genome by traditional short-read sequencing. With the design of the 850K and 50K arrays, this is now possible through high-density SNP marker genotyping, reducing bioinformatic requirements for octoploid strawberry research. In addition to expanding and validating the current molecular toolset, we have demonstrated that allopolyploid reference genomes facilitate the use of straightforward diploid approaches for genomic analysis and quantitative genetics of octoploid strawberry. In doing so, the results of this study help pave the way for molecular breeding of a historically difficult plant genome.

## Materials & Methods

### WGS Sequence Datasets

We generated Illumina sequencing libraries for a diversity panel of 84 wild and domesticated octoploid genotypes (PE150), and the Camarosa reference genotype (PE250). Eight sequenced octoploid libraries (PE100) were obtained from the NCBI sequence read archive (SRA) (SRR1513906, SRR1513893, SRR1513905, SRR1513903, SRR1513892, SRR1513904, SRR1513867, SRR1513873), providing a total of 93 sample libraries in the diversity panel. We generated additional Illumina libraries (PE150) for 189 progeny of an *F.* × *ananassa* x *F. chiloensis* subsp. *lucida* mapping population (Camarosa x Del Norte). Genomic DNA was extracted from immature leaf tissue using the E-Z 96 Plant DNA kit (Omega Bio-Tek, Norcross, GA, USA) with Proteinase K was added to the initial buffer, and RNase treatment delayed until lysate was removed from the cellular debris. An additional spin was added, and incubation steps were heated to 65 C during elution. Libraries were prepared using the KAPA Hyper Plus kit using BIOO Nextflex adapters. DNA was sheared using the Covaris E220 and size selected for an average insert size of 300-nt using magnetic beads (Mag-Bind® RxnPure Plus, Omega Bio-tek). Libraries were pooled and sequenced on the Novaseq at UCSF for average 8x genome coverage in the diversity panel, and 4-8x coverage in the mapping population. DNA capture-based Illumina sequences for the *F. chiloensis* subsp. *lucida* x *F. chiloensis* subsp. *pacifica* population (GP33 x SAL3: n = 46) described in Tennessen et al. (2014), and the *F. virginiana* subsp. *platypetala* x *F. virginiana* subsp. *platypetala* population (KB3 x KB11: n = 46) described in Tennessen et al. (2018) were downloaded from the NCBI SRA.

### Subgenomic WGS Variant Calling

We predicted SNP and indel variants in the Camarosa v1.0 subgenomes using sequenced Illumina short-read libraries for the octoploid diversity panel, Camarosa x Del Norte bi-parental population, and DNA capture sequences downloaded from the SRA. Short-read sequences were quality-trimmed with CutAdapt v1.8 using default parameters and a minimum Phred score of 25. Trimmed short-reads were aligned to the Camarosa v1.0 genome assembly (Edger et al., 2019b) using BWA-mem v0.7.16, processed to mark optical and PCR duplicates using Picard Tools v2.18, and indel-realigned using GATK v3.8. Genomic variants were predicted based on uniquely mapped reads (MapQ > 20) using FreeBayes v1.2 and filtered with vcflib. For analysis of subgenomic variation, a set of hard-filters was applied to remove variants with low site quality (vcflib: QUAL > 40), low contribution of allele observations to site quality (vcflib: QUAL / AO > 10), low read coverage (vcflib: DP > 500), strand bias (vcflib: SAF > 0 & SAR > 0), read-placement bias (RPR > 1 & RPL > 1), unbalanced mapping quality of reference and alternate alleles (vcflib: 0.4 ≤ [MQM / MQMR] ≤ 2.5), unbalanced allele frequencies at heterozygous sites (vcflib: 0.2 ≤ AB ≤ 0.8), low end-placement probability score (EPP ≥ 3), and low strand-bias probability score (vcflib: SRP ≥ 3 & SAP ≥ 3). Sample genotypes were required to have individual read coverage ≥ 4, and at least two reads and a minimum of 0.20 read observations supporting each allele.

### Octoploid Genomic Diversity

We estimated octoploid genetic diversity metrics from a VCF file containing genotype calls for 45M filtered subgenomic SNPs and indels. We calculated the population-level diversity (π) and per-sample heterozygosity of sequence variants using a custom perl script. Chromosomal and genome-wide population nucleotide diversity estimates were calculated as the sum of pairwise diversity for all variant sites divided by total non-gap (N) genomic nucleotides. Sample heterozygosity was calculated as the sum of heterozygous variant sites divided by total non-gap (N) genomic nucleotides, and the fraction of total variant sites that were heterozygous.

### Array Design and Genotyping

Unfiltered genomic variants were filtered to retain sites segregating in *F.* × *ananassa* cultivars. Cultivar variants were filtered to retain biallelic SNP sites with minor allele frequency ≥ 0.05, marker diversity ≥ 0.05, variant QUAL score > 20, and missing data < 15%. Variants requiring 2-probe assays (A/T or C/G) were excluded. 71-nt marker probe sequences were obtained by retrieving 35-nt SNP flanking sequences from the Camarosa v1.0 assembly. Markers containing ambiguous sequences (Ns), or identical probes were excluded. A set of 6.6M probes was submitted to Affymetrix for scoring and recommendation based on strand, kmer uniqueness, and buildability. Probes were scored for likelihood of binding interference or non-specific binding based on off-target variant counts in the binding region, the sum of minor allele frequencies of interfering variants, and counting off-target BLAST alignments (>90% id, >90% query length) in the genome. A final screening panel of 850,000 markers, including 16,000 iStraw probes (Bassil et al., 2015; Verma et al., 2016), was submitted to Affymetrix for constructing the 850K screening array. A panel of 384 octoploid strawberry genotypes was screened on the 850K array. Marker genotype clusters were scored using the Axiom Analysis Suite. Clustering was performed in “polyploid” mode with a marker call-rate threshold of 0.89. Samples were filtered with a dQC threshold of 0.82 and QC CR threshold of 93. A subset of 49,483 probes was selected from polymorphic, QC-passing markers (“PolyHighResolution”, “NoMinorHomozygote”, “OffTargetVariant”) on the 850K screening array to populate the 50K production array. 5,809 LD-pruned (r^2^ < 0.50) probes were pre-selected from the iStraw design, in addition to 47 probes associated with QTL for *Fusarium oxysporum* resistance and the Wasatch day neutral flowering locus (unpublished data). We assigned two markers per gene to a set of 2,878 genes located in expression networks related to flowering and fruit development (Hollender et al., 2014; Kang et al., 2013), or associated with R-gene domains. Non-overlapping 50 kb physical windows were parsed to select single markers containing the highest pairwise diversity in *F.* × *ananassa* genotypes. The remainder of the 50K array was populated by iteratively parsing 50 kb physical windows to select random QC-passing markers for uniform genomic distribution. 1,421 octoploid samples, including the Camarosa x Del Norte mapping population (n = 182), and PI552277 x PI612493 mapping population (n = 96), were genotyped on the 50K array and processed using the Axiom Analysis Suite using the same settings as the 850K dataset.

### WGS Haplotype Linkage Mapping

We used 1.6M female parent informative variant calls (AB x AA), and 1.9M male parent informative variant calls (AA x AB) to generate haploblock markers for mapping genome-wide variant calls in the Camarosa x Del Norte bi-parental population. Camarosa-informative and Del Norte-informative variant calls were divided into parent-specific marker sets, then split by chromosome. For each chromosome, we performed pairwise linkage disequilibrium (LD)-clustering of markers (LD ≥ 0.96) in an initial seed region containing the first 250 chromosome variants, to identify marker groups called in the same phase relative to the unphased Camarosa v1.0 genome. The genotype phase of the LD cluster containing the largest number of markers were selected as the “seed phase”. A 50 kb sliding window was initiated in the seed region and moved across the chromosome, identifying downstream markers in negative LD with the seed phase and reversing the repulsion genotype calls, in order to phase the chromosome into an artificial backcross. If a phasing window skipped a physical region larger than 100 kb without markers, reached a window with fewer than 25 markers, or the average downstream marker LD fell below 0.90, the chromosome was then fragmented at the breakpoint, seed-phase clustering repeated, and the sliding window reset for the subsequent downstream region. We used the software PhaseLD (Marand et al., 2017) with a 50-marker window and 1-marker step size to predict crossover events in the backcross-phased chromosome blocks. Window-specific variant calls lying between the predicted recombination breakpoints were used to generate consensus genotypes representing the haploblock. We mapped the haploblock markers using software ONEMAP (Margarido et al., 2007) and BatchMap (Schiffthaler et al., 2017) in outcross mode. ONEMAP was used to bin co-segregating markers, calculate pairwise recombination fractions, determine optimal LOD thresholds, then cluster markers into linkage groups based on a LOD threshold of 8, and maximum recombination fraction of 0.22. Marker orders and genetic distances were re-estimated in parallel with BatchMap using a window of 50 markers, window overlap of 30, and ripple window of 6 markers.

### Array and DNA Capture Linkage Mapping

We performed single-marker linkage mapping of populations genotyped using the 50K array or DNA capture sequences because each contained fewer than 10,000 segregating markers per parent. Individual parent genotypes were mapped separately using their respective informative marker subsets. We filtered markers based on a chi-square test for segregation distortion (p-value < 0.10), and excluded markers with >5% missing data. ONEMAP was used to bin co-segregating markers, calculate pairwise recombination fractions, determine optimal LOD thresholds, and cluster markers into linkage groups based on a LOD threshold of 8, and maximum recombination fraction of 0.22. Marker orders and genetic distances were re-estimated in parallel with BatchMap using a window of 20 markers, window overlap of 15, and ripple window of 6 markers.

### GDR Microsatellite Alignment

We obtained the complete set of microsatellite primers developed in *Fragaria* species from the Rosaceae Genomics Database (GDR). Primers pairs were aligned to the Camarosa v1.0 genome in an orientation-aware manner using IPCRESS with a maximum amplicon fragment size of 500 bp and allowing 1 mismatch per primer.

## Author Contributions

MAH, PPE, and SJK contributed conception and design of the study; MAH, AL, and MJF performed the statistical analysis; RF, CA, and GC provided genetic material and collected, submitted DNA samples; MAH and SJK wrote the first draft of the manuscript; All authors contributed to manuscript revision, read and approved the submitted version.

## Acknowledgments

We thank our collaborators at Affymetrix for constructing the 50K and 850K octoploid strawberry SNP genotyping arrays.

## Data Availability

The datasets generated for this study can be found in the NCBI Sequence Read Archive (https://www.ncbi.nlm.nih.gov/sra).

