## Supplemental Materials for "Chromosome Evolution of Octoploid Strawberry"

**Supplementary Material**

Supplementary datasets, figures and tables.

**Supplementary Data Files**

**Supplementary Dataset 1.** Summary of octoploid genetic maps generated by WGS sequencing, 50K SNP array, and DNA capture datasets.

**Supplementary Dataset 2**. 850K screening array diversity panel of octoploid strawberry cultivars and wild accessions.

**Supplementary Dataset 3.** Panel of 446,644 validated marker probes from the 850K SNP screening array.

**Supplementary Dataset 4.** Panel of 49,483 marker probes selected for populating the 50K SNP production array.

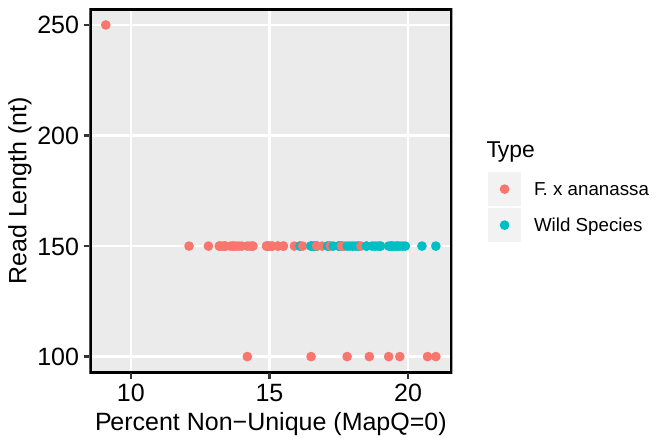

**Supplementary Figure 1.** Frequency of non-unique short-read alignments from diversity panel WGS libraries aligned to the Camarosa v1.0 genome assembly.

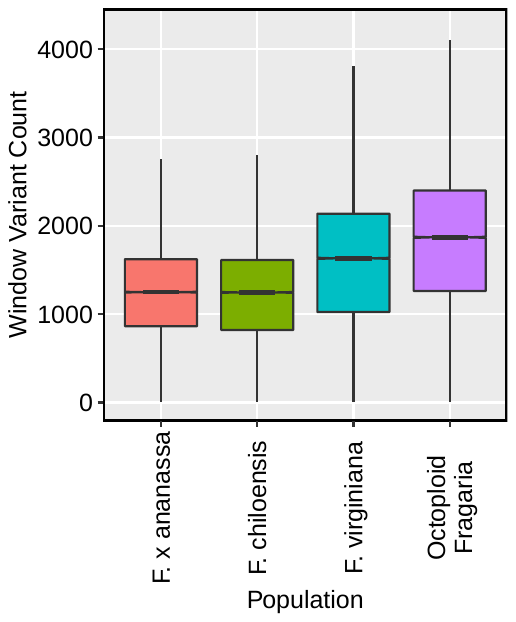

**Supplementary Figure 2.** Distribution of total subgenome-specific variant calls in 50 kb physical windows spanning the Camarosa v1.0 genome assembly, including domesticated samples (*F.* × *ananassa*), both wild progenitor species (*F. chiloensis*, *F. virginiana*), and all octoploid samples (Octoploid *Fragaria*)

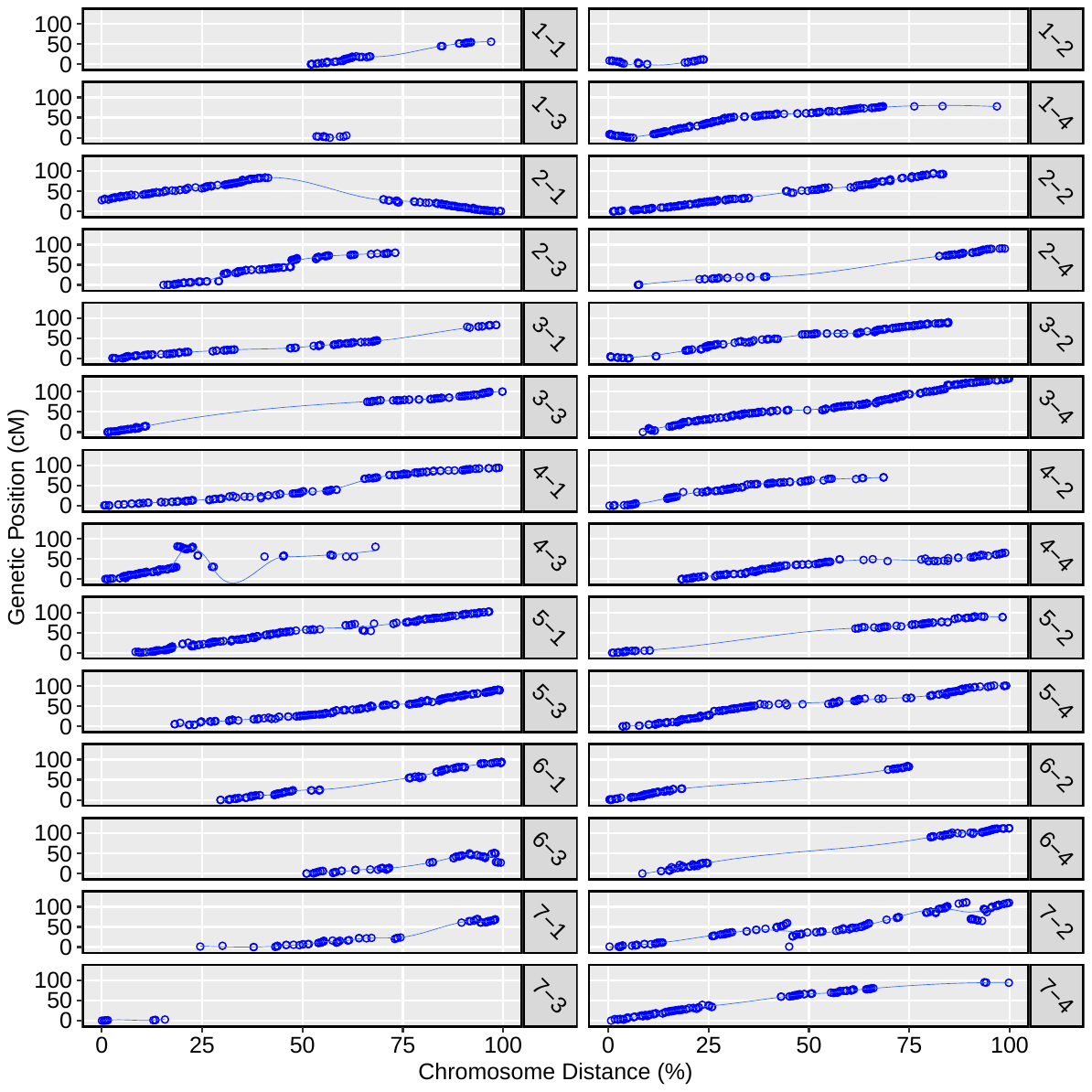

**Supplementary Figure 3.** Haplotype map of commercial *F.* × *ananassa* cultivar Camarosa plotted against the Camarosa v1.0 physical genome.

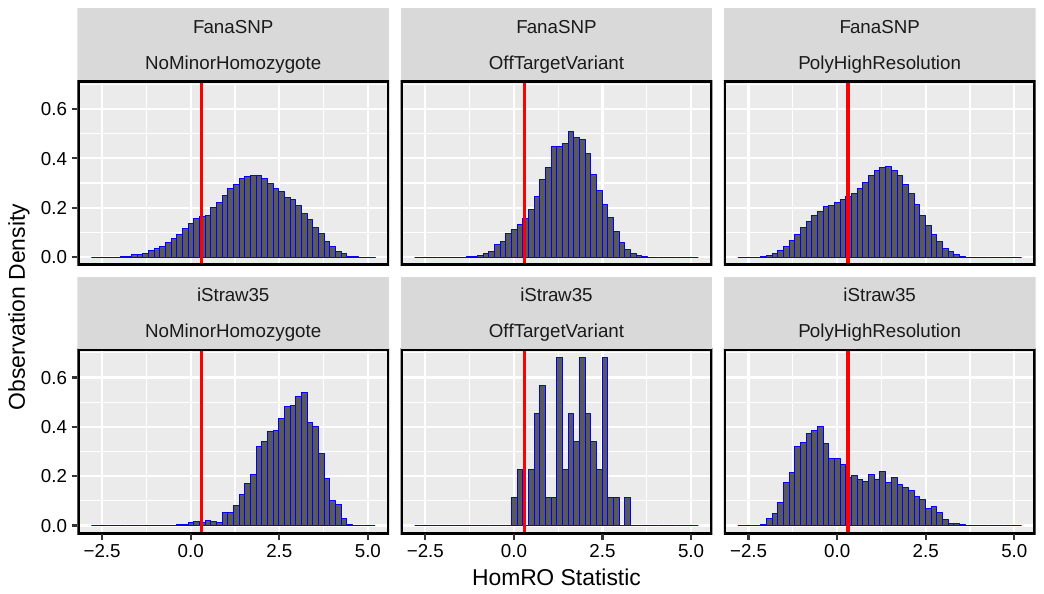

**Supplementary Figure 4.** Distribution of HomRO statistic for 446,644 validated marker probes. Distributions are compared between probes designed in the current study (FanaSNP), and legacy probes retained from the iStraw35 SNP array, in three polymorphic marker classes: PolyHighResolution, NoMinorHomozygote, and OffTargetVariant. Vertical red line indicates threshold for copy-specific probe binding.

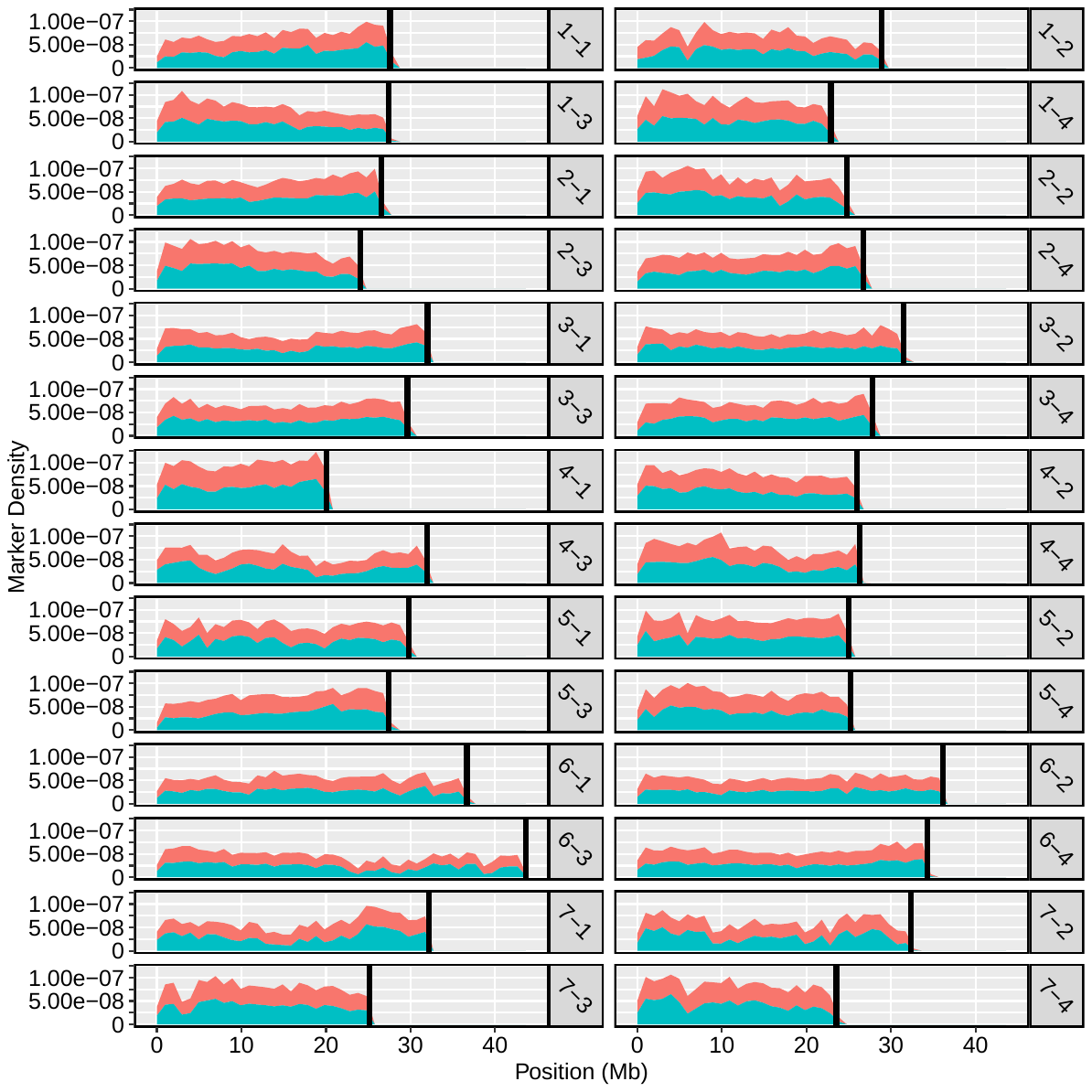

**Supplementary Figure 5.** Chromosome-wide marker density showing the distributions of total 850K array markers (blue) and total 50K array markers (red) in the Camarosa v1.0 genome.

**Supplementary Figure 6.** Comparative mapping of four wild octoploid subspecies (*F. chiloensis* subsp. *lucida*, *F. chiloensis* subsp. *pacifica*, *F. virginiana* subsp. *platypetala*, *F. virginiana* subsp. *virginiana*) across the 28 octoploid strawberry chromosomes.

**Supplementary Table 1.** WGS diversity panel of octoploid strawberry cultivars and wild accessions.

| **Sample Name** | **ID/Accession** | **Species** |  | 2 Futalefu 6A introgressed | PI616554 | F. chiloensis subsp. chiloensis forma chiloensis |
| --- | --- | --- | --- | --- | --- | --- |
| 04C076P004 | 04C076P004 | F. × ananassa |  | Ambato ecuador | PI616766 | F. chiloensis subsp. chiloensis forma chiloensis |
| 08C123P001 | 08C123P001 | F. × ananassa |  | Ambato peru | PI551736 | F. chiloensis subsp. chiloensis forma chiloensis |
| 10C144P002 | 10C144P002 | F. × ananassa |  | CFRA 1377 | PI616683 | F. chiloensis subsp. chiloensis forma chiloensis |
| 11C036P601 | 11C036P601 | F. × ananassa |  | Isle of Lemuy 02A White | PI552038 | F. chiloensis subsp. chiloensis forma chiloensis |
| 11C141P001 | 11C141P001 | F. × ananassa |  | 2 TAP 4B | PI612317 | F. chiloensis subsp. chiloensis forma patagonica |
| 11C153P003 | 11C153P003 | F. × ananassa |  | CFRA 1064 | PI616516 | F. chiloensis subsp. chiloensis forma patagonica |
| 65C065P601 | 65C065P601 | F. × ananassa |  | CFRA 1065 | PI616517 | F. chiloensis subsp. chiloensis forma patagonica |
| 71C098P605 | 71C098P605 | F. × ananassa |  | PI552058 | PI552058 | F. chiloensis subsp. chiloensis forma patagonica |
| 94C016P001 | 94C016P001 | F. × ananassa |  | PI552092 | PI552092 | F. chiloensis subsp. chiloensis forma patagonica |
| Aberdeen | PI551630 | F. × ananassa |  | PI602569 | PI602569 | F. chiloensis subsp. chiloensis forma patagonica |
| Albion | 97C117P003 | F. × ananassa |  | PI616531 | PI616531 | F. chiloensis subsp. chiloensis forma patagonica |
| Cabrillo | 08C181P001 | F. × ananassa |  | San Martin | PI552295 | F. chiloensis subsp. chiloensis forma patagonica |
| Camarosa | 88C029P603 | F. × ananassa |  | CA 1405 | PI551731 | F. chiloensis subsp. lucida |
| Diamante | 91C248P006 | F. × ananassa |  | Del Norte | PI551753 | F. chiloensis subsp. lucida |
| Douglas | 70C003P108 | F. × ananassa |  | Honeyman Par | PI612489 | F. chiloensis subsp. lucida |
| Dover x Cam F2-34 | NA | F. × ananassa |  | LCM-10 | PI551468 | F. chiloensis subsp. lucida |
| Earliglow | PI551394 | F. × ananassa |  | BSP-03 | PI551467 | F. chiloensis subsp. pacifica |
| Emily | PI616854 | F. × ananassa |  | CA 1499 | PI551735 | F. chiloensis subsp. pacifica |
| Ettersburg121 | PI551904 | F. × ananassa |  | Hartney Bay | PI616770 | F. chiloensis subsp. pacifica |
| Fenella | NA | F. × ananassa |  | KH 94-06 | PI616652 | F. chiloensis subsp. pacifica |
| FL_13.26.134 | NA | F. × ananassa |  | LPB 2-01 | PI551458 | F. chiloensis subsp. pacifica |
| FL_13.55.195 | NA | F. × ananassa |  | Scotts Creek | PI612490 | F. chiloensis subsp. pacifica |
| FL_14.100.59 | NA | F. × ananassa |  | Yaquina A | PI551765 | F. chiloensis subsp. pacifica |
| Florida Elyana | NA | F. × ananassa |  | BC6 | PI660767 | F. virginiana subsp. glauca |
| Florida127 | NA | F. × ananassa |  | CA 1226 Big Cottonwood | PI612491 | F. virginiana subsp. glauca |
| Fronteras | 08C132P608 | F. × ananassa |  | CFRA 420 | PI551775 | F. virginiana subsp. glauca |
| Gorella | PI551483 | F. × ananassa |  | LH 18-2 | PI551876 | F. virginiana subsp. glauca |
| Guardian | PI551407 | F. × ananassa |  | LH 20-1 | PI551877 | F. virginiana subsp. glauca |
| Holiday | PI551653 | F. × ananassa |  | RH 43 Delta Junction | PI612496 | F. virginiana subsp. glauca |
| Hood | PI551502 | F. × ananassa |  | JP 95-1-1 McLellan | PI612570 | F. virginiana subsp. grayana |
| Howard 17 | PI551593 | F. × ananassa |  | NC 95-19-1 | PI612486 | F. virginiana subsp. grayana |
| Jucunda | PI551623 | F. × ananassa |  | IH-35 | PI551470 | F. virginiana subsp. platypetala |
| Kaiser's Samling | PI270471 | F. × ananassa |  | Steven's Pass | 15X001P001 | F. virginiana subsp. platypetala |
| Korona | PI551581 | F. × ananassa |  | Straw Mt | PI616601 | F. virginiana subsp. platypetala |
| Mara De Bois | 17Z001P001 | F. × ananassa |  | CFRA 1927 | PI657849 | F. virginiana subsp. virginiana |
| Palomar | 00C259P002 | F. × ananassa |  | Frederick 9 | PI612493 | F. virginiana subsp. virginiana |
| Pelican | PI637960 | F. × ananassa |  | Hinesburg | PI552277 | F. virginiana subsp. virginiana |
| Portola | 01C206P005 | F. × ananassa |  | KY-08 | PI616568 | F. virginiana subsp. virginiana |
| Puget Reliance | PI664321 | F. × ananassa |  | LH 50-4 | PI612495 | F. virginiana subsp. virginiana |
| Reikou | PI616627 | F. × ananassa |  | N-8 | PI616676 | F. virginiana subsp. virginiana |
| Seascape | 83C049P001 | F. × ananassa |  | NC 95-11-1 | PI616691 | F. virginiana subsp. virginiana |
| Senga Sengana | PI264680 | F. × ananassa |  | NC 96-25-1 | PI616800 | F. virginiana subsp. virginiana |
| Sweet Charlie | PI664317 | F. × ananassa |  | NC 96-5-3 Chadwick | PI612325 | F. virginiana subsp. virginiana |
| Tioga | 53C009P002 | F. × ananassa |  | Sheldon | PI551651 | F. virginiana subsp. virginiana |
| White Carolina | PI551681 | F. × ananassa |  | UC-11 | PI551495 | F. virginiana subsp. virginiana |
| Wiltguard | 52C016P007 | F. × ananassa |  |  |  |  |
| Winter Dawn | none | F. × ananassa |  |  |  |  |

**Supplementary Table 2.** Genome-wide summary of population-specific and overall octoploid strawberry nucleotide diversity (π) across the 28 ancestral diploid chromosomes.

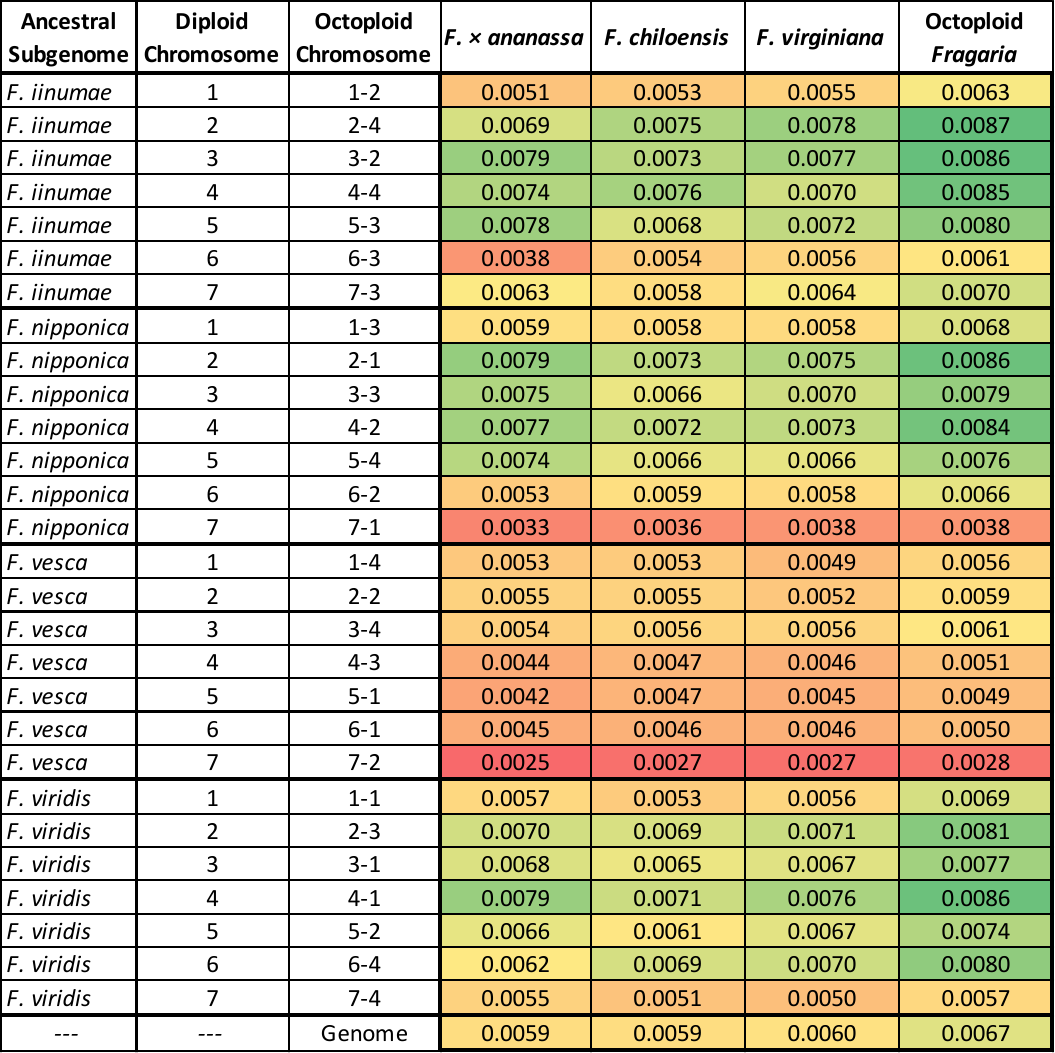

**Supplementary Table 3.** Summary of octoploid genetic mapping results based on 50K SNP array genotyping, including per-chromosome marker densities (SNP/Mb), numbers of unique and co-segregating variant sites, and map sizes (cM).

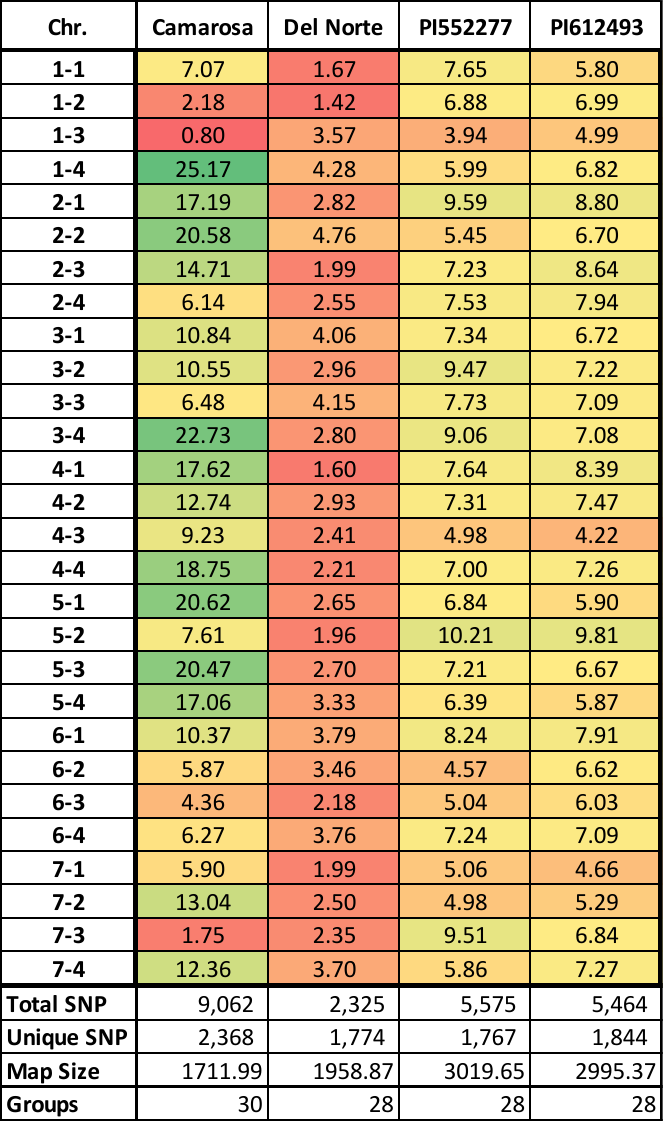
